# Optimal Sampling Rate for 3D Single Molecule Localization

**DOI:** 10.1101/2023.09.19.558416

**Authors:** Huanzhi Chang, Shuang Fu, Yiming Li

**Affiliations:** Department of Biomedical Engineering, Guangdong Provincial Key Laboratory of Advanced Biomaterials, Southern University of Science and Technology, Shenzhen, China

## Abstract

Resolution of single molecule localization microscopy (SMLM) depends on the localization accuracy, which can be improved by utilizing engineered point spread functions (PSF) with delicate shapes. However, the intrinsic pixelation effect of the detector sensor will deteriorate PSFs under different sampling rates. The influence of the pixelation effect to the achieved 3D localization accuracy for different PSF shapes under different signal to background ratio (SBR) and pixel dependent readout noise has not been investigated in detail so far. In this work, we proposed a framework to characterize the 3D localization accuracy of pixelated PSF at different sampling rates. Four different PSFs (astigmatic PSF, double helix (DH) PSF, Tetrapod PSF and 4Pi PSF) were evaluated and the pixel size with optimal 3D localization performance were derived. This work provides a theoretical guide for the optimal design of sampling rate for 3D super resolution imaging.

## 1. Introduction

The improvement of imaging resolution had always been taken top priority during the development of microscopy techniques. It was first theoretically derived by Abbe in 1873 that the resolution is confined by ∼λ⁄2NA (NA is the numerical aperture of the objective), known as the diffraction limit. The development of super-resolution microscopy has surpassed this limit and brought optical microscopy into the era of nanoscopy, enabling resolving biological structures at the molecular scale. Single-molecule localization microscopy (SMLM), such as PALM [1, 2] and STORM [3], is one of the most widely used super-resolution techniques. Basic procedure in SMLM contains isolation of individual emitters and precise localization of diffraction limited single molecule images. Despite different acronyms, PALM/STORM share the same requirement in separating emitters: Only one of them is allowed to fluoresce, whereas its neighbors have to remain silent [4]. While this sequential on-and-off switching of molecular fluorescence is highly effective at making neighboring molecules discernible, it doesn’t provide precise location. Precise localization is normally achieved by fitting measured images to a predefined PSF model.

One of the most important questions in SMLM is the achievable localization accuracy of single molecule images. In general, it is affected by several factors, such as PSF model, SBR, camera noise and pixelation effect. Upon proper modeling, it has been shown that maximum likelihood estimation could achieve theoretical minimum uncertainty of single molecule localization in simulated data (Cramer-Rao lower bound, CRLB) [5]. Recently, by exploiting spline interpolated experimental PSF model, we have shown that CRLB could be achieved even in real experimental conditions [6].

Most of previous works tried to improve the localization accuracy by PSF engineering or optimal localization algorithms. Few of them systematically considered the effect of the sampling rate on the achievable localization accuracy, commonly denoted by CRLB. Therefore, it is still not clear that the achieved localization accuracy is with an optimal sampling rate (OSR) using the pixelated 2D array detectors. Normally, larger pixel sizes result in a deteriorated localization precision since the locations of all the photons within a single pixel is unknown. On the other hand, the decrease of the pixel size will also result in more pixels within a certain area of the single molecule pattern. Therefore, more detector noise will be introduced. Besides, the tradeoff between pixel size and field of view (FOV) makes it equally significant to consider a PSF’s performance under low sampling rates when large FOV imaging is performed using cameras with limited pixels. One of the most widely accepted theories for the proper pixel size selection for single molecule localization is based on the work by Thompson et al. in 2002[7]. They pointed out that the optimal pixel size should be about equal to the standard deviation of the point spread function. However, the theory is based on a Gaussian PSF model and only lateral localization precision was considered. Other works suggested to use a more precise Airy PSF model and MLE based single molecule localization [8, 9] to achieve the CRLB determined by information theory. Still, only lateral localization precision was taken account for, and the readout noise is uniform across pixels which is only valid for CCD cameras.

In the past ∼15 years, various 3D SMLM techniques were developed, allowing accurate analysis of subcellular structures in all three dimensions. Conventional Gaussian PSF is no longer suitable for this purpose due to its symmetry between both sides of the focal plane. By encoding the z positions to the shape of the PSF, PSFs with different shapes were generated, such as astigmatic PSF [10], DH-PSF [11-13], Tetrapod PSF [14], and 4Pi PSF [15-17]. These methods break the symmetrical shape of light fields and conserve the energy during defocus in manifold forms. For example, the Tetrapod PSF has two concentrated sidelobes with different spreading directions above and below the focal plane. Its effective depth of field (DOF) can even reach 20 *μ*m, which is much larger than the DOF of a high NA objective[18]. Since z positions are determined by the shape of the PSFs, the sampling rate will undoubtedly affect the z localization precision. However, it has not been systematically investigated so far. Moreover, sCMOS camera has become more and more popular in scientific imaging for its fast imaging speed, large dynamic range, and ultra-high pixel number. However, its readout noise is often pixel dependent. To date, the impact of the pixelation effect on the 3D localization accuracy of PSFs with different shapes under pixel dependent readout noise has not been quantified or explored in detail.

Here, we established a framework to simulate the pixelated PSFs with different sampling rate using the vectorial PSF model. In this framework, we first generated a 3D PSF model with fine sampling rate (2 nm) and binned it to large pixels to model the pixelated PSF with large sampling rate. We then quantitatively evaluated the pixelation effect on the localization accuracy of four different kinds of PSFs (astigmatic PSF, DH-PSF, Tetrapod PSF and 4Pi PSF). The optimal pixel sizes to achieve the best 3D CRLB under different SBRs for each PSF were obtained. Besides, we also explored the localization bias due to the pixel dependent readout noise under different sampling rates.

## 2. Methods

### 2.1 Pixelated Vector PSF simulation

To accurately simulate the PSF of a high NA objective which is commonly used in superresolution imaging, a vectorial PSF model that accounts for the light polarization and Fresnel transmission coefficients under high angle refraction is used in this work [19-21]. Since most of the fluorescent probes often rotate freely, we modeled the PSF as the summed image of three orthogonal dipoles with equal strength to mimic the random orientation case. As shown in Fig. 1a, the PSF calculation can be divided into two steps: 1) First we calculate the basic light field components 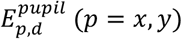 at the pupil plane contributed by each dipole component *d* = *x, y, z* at the object plane, respectively. The electrical field at the pupil is further modified both in amplitude and phase to take account for the energy change and aberrations during the light transmission. 2) The light field at the pupil plane is then formed into an image on the camera by the tube lens. Since all ray angles passing through the tube lens are normally small, scalar diffraction theory are applied:

**Fig. 1.**
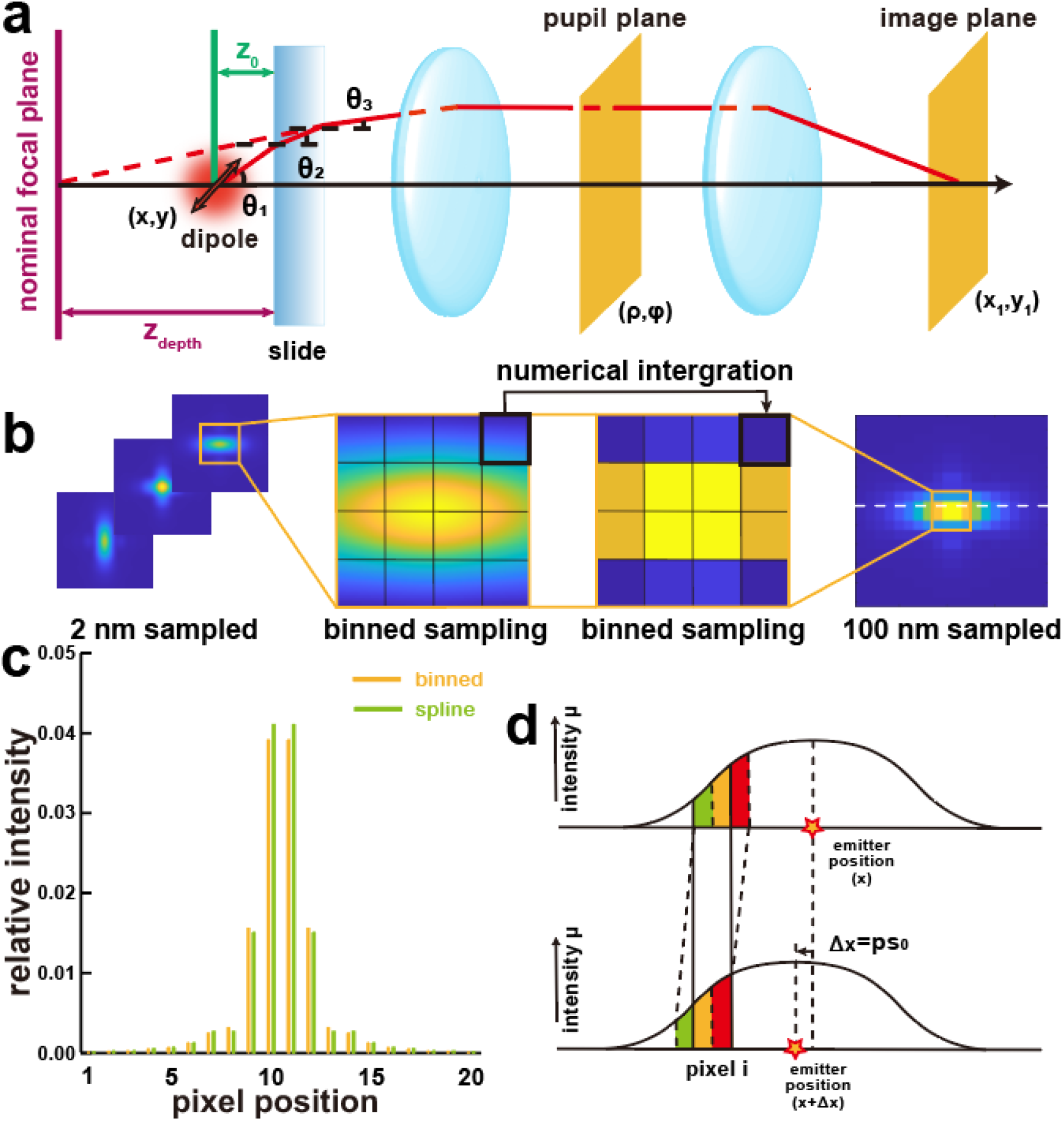
Schematic of PSF simulation and derivative computation. (a) Vector PSF simulation model. Propagation from dipole emitter to pupil plane describes 3 orthogonal electric field components contributing to p and s polarization components at pupil plane. Propagation from pupil plane to image plane can be implemented by Fourier transform of the pupil function. The Fourier transform was implemented using Bluestein’s algorithm. By inserting different pupil phases at pupil plane, we can simulate different PSFs. (b) Illustration of binned astigmatic PSF generation. Firstly, a 2 nm fine sampled PSF size of 500×500 was simulated. By combing nearby 50×50 sub-pixels into a large pixel, a 100 nm sampled PSF size of 20×20 was simulated. (c) Comparation of binned PSF and spline interpolated PSF. Line plots as indicated in b. (d) 1D-illustration of computation method for derivative with respect to emitter position 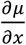 used in CRLB calculation. Green area minus red area denotes the intensity change in pixel i when moving the emitter Δ*x* left.

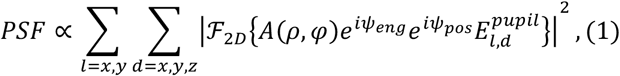

where *ℱ*_*2D*_ denotes the 2D Fourier transform. *A*(*ρ, φ*) = (1 − *ρ*^*2*^*NA*^*2*^*/n*_1_^*2*^)^−¼^ is an amplitude function considering aplanatic correction factor. (*ρ, φ*) is the normalized polar coordinate in the pupil plane with *ρ* = 1 corresponding to the limiting aperture angle *NA/n*_3_. *n*_1_, *n*_*2*_ and *n*_3_ is the refractive index of sample, cover slip and immersion oil, respectively. *ψ*_*eng*_ is the modulated phase for PSF engineering and *ψ*_*pos*_ is the emitter position dependent phase shift:

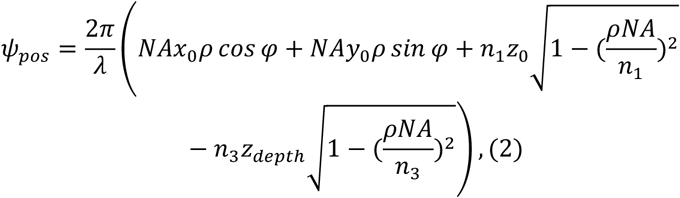

where λ is the wavelength, (*x*_0_, *y*_0_, *z*_0_) is the emitter position. *z*_0_ represents the emitter’s depth away from the cover glass. *z*_*depth*_ is the distance between the nominal focal plane and the cover glass. The basic light field components 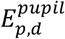 at the pupil plane are given by:

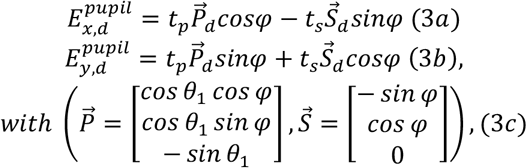

where 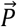 and 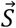 are the basis polarization vectors. *t*_*p*_ and *t*_*s*_ are the total Fresnel transmission coefficients for the p- and s-polarized light through multiple mediums: *t*_*k*=*p,s*_ = *t*_*k*,1−*2*_ × *t*_*k,2*−3_. *θ* and *φ* are the polar and azimuthal angles respectively. Take the boundary 1 − *2* as an example, the Fresnel transmission coefficients are given by:

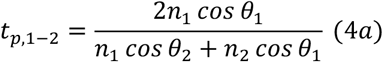

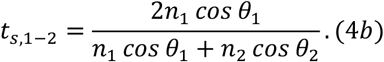

To ensure an accurate result of the numerical implementation, the minimum number of sampling points *M* on 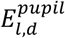 should satisfy the below relationship according to [22]:

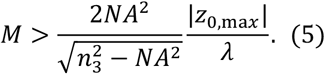

In this work, we assume a refractive index match case with the following parameter settings: *NA* = 1.49, *n*_1,*2*,3_ = 1.518, λ = 660 *nm*, and |*z*_0,max_| = 1500 *nm* (maximum defocus distance of PSFs quantified in this work). These require *M* > 35 and we set it as 128 for improving accuracy. Chirp-z transformation was used to perform the Fourier transform between pupil plane and image plane since it is flexible to calculate the light field with different sampling rate[22].

### 2.2 Noise simulation

After building an ideal vector PSF model in the previous section, noise is then added to generate the experimental pixelated PSF images:

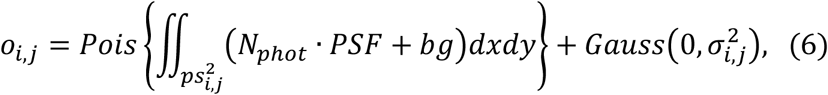

where *o*_*i,j*_ is the observation at pixel (*i, j*), Pois(x) is a Poisson random number with mean value of x and *Gauss*(*u*, σ^*2*^) is a Gaussian random number with mean value of *μ* and standard deviation of σ. Four types of noise are considered: 1) *bg* is the background noise arises from out of focus fluorescence signal, in the unit of photoelectrons/nm^2^; 2) Pixelation noise denotes the continuous *PSF* model is integrated over each pixel (*i, j*) with size of *ps* nm; 3) Shot noise arises from photon collecting procedure. Due to the discrete nature of photon emission, the number of photons collected by camera detector in a certain area within a certain time follows the Poisson distribution; 4) Readout noise arises from the electron/voltage conversion process that can be described by a Gaussian distribution. The readout noise (σ_*i,j*_) 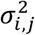 is pixel-dependent for sCMOS camera, while it is uniform for CCD camera. *N*_*phot*_ is the number of photoelectrons. We omit deterministic parameters that will not change the distribution property such as quantum efficiency, analog-to-digital conversion factor and camera baseline, etc. The expected value *μ*_*i,j*_ at the pixel (*i, j*) can be given by:

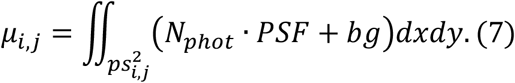

The direct computation of the integral above is challenging as all data is discrete in silico. Here, we first simulated a fine-sampled vector PSF with tiny pixel size *ps*_0_, then binned several tiny pixels into a large pixel to get the PSF image with desired pixel size *ps* to mimic the pixelation effect (Fig. 1b). The number of tiny pixels used for each large pixel is 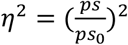. This process is denoted as pixel-binning simulation in the following paragraph and can be described as below:

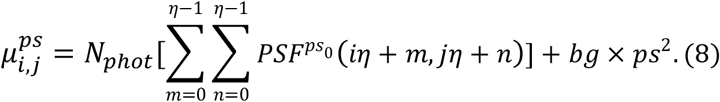

In this work the fine-sampled model 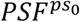 was simulated with *ps*_0_ = *2 nm* over a 6000 × 6000 *nm*^*2*^ region with (*m, n*) the grid index. z step is dependent on the DOF for each type of engineered PSF due to memory limit. Take the astigmatism PSF with *ps* = 100 *nm* as an example, *η*^*2*^ = 2500 tiny pixels are added to get a large pixel (Fig. 1b). As shown in Fig. 1c, the difference between the binned PSF model *μ*^100 *nm*^ and the spline interpolated *PSF*^100 *nm*^ is obvious.

### 2.3 Localization accuracy

CRLB indicates the minimum variance that an unbiased parameter estimator can reach. In SMLM, it is widely used to evaluate the localization performance of different PSF models [5, 8, 23]. Therefore, we calculated CRLB of different engineered PSF models under different pixel sizes to find the optimal spatial sampling rate. The CRLB is given by the inverse of the Fisher information matrix:

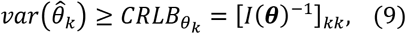

where ***θ*** = [*x, y, z, N*_*phot*_, *b*] is the set of parameters to be estimated, including emitter position, photon, and background. *I*(***θ***) is the Fisher information matrix described as:

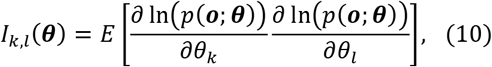

with *p*(***o; θ***) the likelihood of the observation ***o*** given the model parameter ***θ***. By approximating the combined distribution of Eq. (6), which contains a Poisson process and a Gaussian process, we arrived a single Poisson distribution according to [24]:

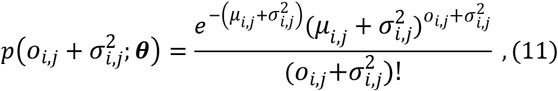

where *μ*_*i,j*_ and σ_*i,j*_ stand for the expected photoelectrons and readout noise at the pixel (*i, j*), respectively. This gives the Fisher information matrix as described similarly in previous publications [24]:

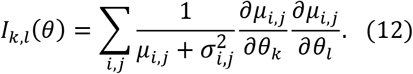

Here, based on the binned PSF model as described above, we used numerical differentiation to calculate the derivative. The partial derivative with respect to x, photons and background using the numerical differentiation can be given by:

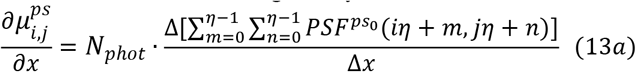

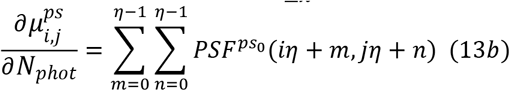

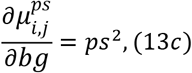

where *ps* is the binned pixel size, 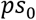 is the fine-sampled PSF with pixel size of *ps*_0_. Thus, each binned pixel contains 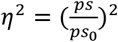 subpixels. We gave a 1D demonstration to show the derivatives with respect to emitter position (Fig. 1d). Given that Δ*x* = *ps*_0_, moving the emitter Δ*x* left is equivalent to moving the sensor’s pixel a grid unit (*ps*_0_) right, and change in *μ*_*i*_ can be expressed as the red part minus green part. When applied to a 2D situation, the derivatives with respect to x can be computed by:

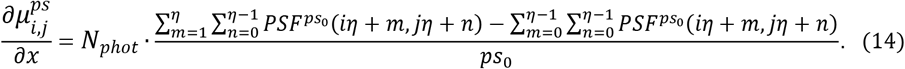

It can be seen that two terms of numerator in Eq. (14) contains much overlap region, thus it can be simplified to:

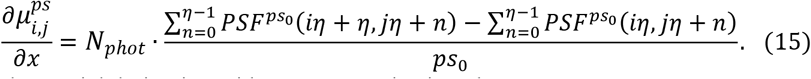

Similarly, partial derivative with respect to *y* is given by:

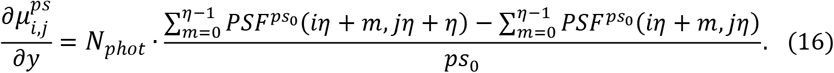

Partial derivative about *z* is calculated by:

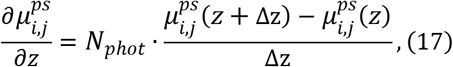

where Δz is the sampling rate in z direction. The CRLB for the estimation of ***θ*** can be achieved according to Eq. (9), and CRLB_3d_ is derived from the sum of the CRLB_x_, CRLB_y_, CRLB_z_:

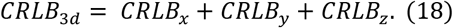

To quantify the CRLB across the whole depth of field for each PSF, we averaged CRLB_3d_ at different z-depth with a fixed sampling interval along optic axis (10 nm for astigmatic PSF, 15 nm for DH PSF, 20 nm for Tetrapod PSF, 10 nm for 4Pi PSF) within the entire z-depth(±600 nm for astigmatic PSF, ±900 nm for DH PSF, ±1500 nm for Tetrapod PSF, ±600 nm for 4Pi PSF). Moreover, the CRLB may vary with a PSF’s center position moving in a fine-sampled grid size and we compensate for this effect by averaging these CRLBs.

## 3. Results

Next, we explored the effects of pixelation on localization precision of four different PSFs (Astigmatic, DH, Tetrapod, and 4Pi PSF) using 3D CRLB. We found that each PSF behaved differently, exhibiting varying 3D CRLB responses to the pixelation effect. To understand the principle behind, we focused on the optimal sampling rate with the lowest 3D CRLB and how the sensitivity of 3D CRLB to sampling rate varies with different PSFs and background conditions. By 1) analyzing 3D CRLB under different SBR and 2) comparing the optimal sampling rate and CRLB sensitivity of different PSFs, we identified the problem as a tradeoff between SBR and the number of delicate features of the PSF. All simulations were conducted with the following parameters, unless specified differently: wavelength λ = 660 *nm*, numerical aperture *NA* = 1.49, refractive index of sample, cover slip and immersion oil *n*_1,*2*,3_ = 1.518, photon number *N*_*phot*_ = *2*000, background photons *bg* = *2* × 10^−3^ *photons/ nm*^*2*^, readout noise at each pixel σ = *2 photo electrons*.

### 3.1 Optimal sampling rate for astigmatic PSF

Astigmatic PSF is the most widely used PSF for 3D imaging as it can be easily implemented by inserting a cylindrical lens in the imaging path. To simulate an astigmatic PSF, we set the astigmatism term of the pupil function as 80 nm (Fig. 1a). The imaging range of astigmatic PSF is normally in the range of 600 nm above and below the focus. The 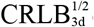 quickly deteriorates beyond this axial range (Fig. 1b). If there is no readout noise, smaller pixel size (finer sampling) will result in better 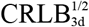. Due to the existence of readout noise, smaller pixel size will introduce more noise for ROIs of same area. In contrast, large pixel size will result in the loss of precise localization of each photon within individual pixels. Therefore, the optimal pixel size should have a best tradeoff between readout noise and sparse-sampling effect. Here, we systematically evaluate the effects of the pixel size on the achieved 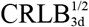 and the optimal pixels size which achieved the best 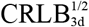 under different SBRs. Considering the precision of the differential method used, the pixel size range evaluated is set between 30 nm and 250 nm. We first explored astigmatic PSF’s 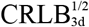 as a function of pixel size at different focal plane. As shown in Fig. 2c, the 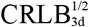 exhibits the same downward-upward trend at all focal plane as the increase of pixel size. Moreover, when focal plane changes from 600 nm to 0 nm, the optimal pixel size decreases from 105 nm to 66 nm. We also found that the sensitivity of 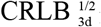 relative to pixel size (defined as the relative increase in 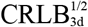 from optimal pixel size to 250nm) is also different at different focal plane. The sensitivity increases from 12.5% to 50.4% from focal plane of 600 nm to 0 nm. It is probably due to the fact that PSF near focus has a sharper pattern, thus becoming more sensitive to sparse sampling (in other words a large pixel size). We then analyzed the effect of the pixel size on 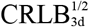 under different SBR (Fig. 2d). When *N*_*phot*_ changes from 2000 to 20000, optimal pixel size decreases from 93 nm to 81 nm, indicating that better SBR has smaller optimal pixel size which introduces more readout noise for same imaging area. For different SBRs, the 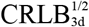 sensitivity to the pixel size does not change much compared to that for PSFs at different focal plane (Fig. 2c and d), suggesting that the shape of the PSF plays a more important role regarding to the change of 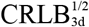 at different pixel size. We then evaluate the pixelation effect on the lateral and axial CRLB separately. When *N*_*phot*_ changes from 2000 to 20000, the optimal pixel size remains almost unchanged (∼100 nm) and the sensitivity slightly decreases from 17.1% to 12.7% for 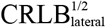 (Fig. 2e), while it decreases from 84 nm to 58 nm and the sensitivity slightly increases from 38.0% to 41.0% for 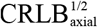, (Fig. 2f). The sensitivity for 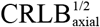 is much higher than 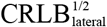 indicating that axial localization accuracy is more vulnerable to pixelation than lateral localization accuracy for astigmatic PSF.

**Fig. 2.**
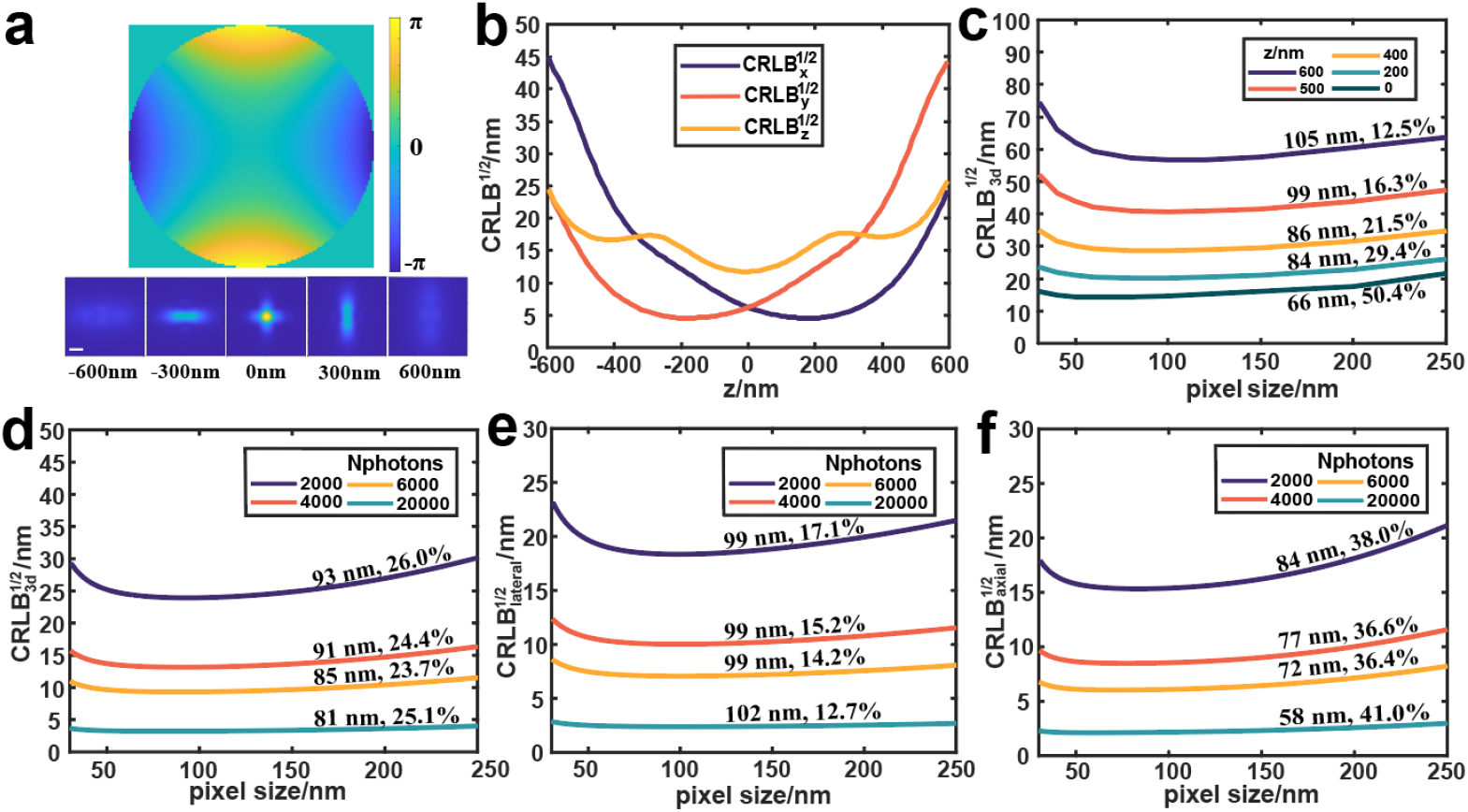
Astigmatic PSF simulation and CRLB calculation. (a) Astigmatic pupil and corresponding PSF. (b) 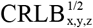 at different focal plane. (c) 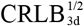 versus pixel size at different focal plane. (d-f) CRLB^½^ versus pixel size under different SBR, for 3d, lateral dimension, axial dimension, respectively. Scale bar 1*μ*m.

### 3.2. Optimal sampling rate for double-helix PSF

DH PSF was named after its “double helix” structure along the axial dimension. It has two lobes, and they spin around their midpoints throughout the ∼2 *μ*m depth of field. The overall size of the DH PSF does not change much within this axial range. Here, rings of azimuthal ramps were used to construct a DH PSF[12, 25]. Pupil function with 3 rings of azimuthal ramps whose increasing slope with a ring radius ratio *α* of 1/2 was used [20]. For a 2*π* rotation, the axial range analyzed was set as ±900 nm (Fig. 3a). We first evaluated the change of 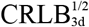 as a function of pixel size at different focal plane. As shown in Fig 3c, a similar downward-upward trend of 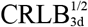 was observed as that of astigmatic PSF. However, both optimal pixel size (∼100 nm) and sensitivity (∼16%) remains almost unchanged at different focal plane for DH-PSF which is different compared to the behavior of astigmatic PSF. This is probably because DH-PSF shows similar patterns which only rotate at different axial positions. Fig. 3d-f shows CRLB versus pixel size under different SBR for 3D, lateral and axial, separately. Compared to astigmatic PSF, DH-PSF has a larger optimal size for 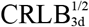 (decreases from 101 nm to 96 nm when the photons increase from 2000 to 20000) and a lower pixelation sensitivity (∼ 17%). When *N*_*phot*_ changes from 2000 to 20000, optimal pixel size for 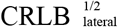 remains about 100 nm, while it changes from 103nm to 90nm for 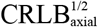 (Fig. 3e, f). Different with astigmatic PSF, the pixelation sensitivity of 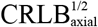 (∼13%, Fig. 3f) is much lower than that of 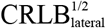 (∼21%, Fig. 3e). It is probably because the axial localization of DH PSF is mainly based on its rotation angle which is pixel-size-insensitive.

**Fig. 3.**
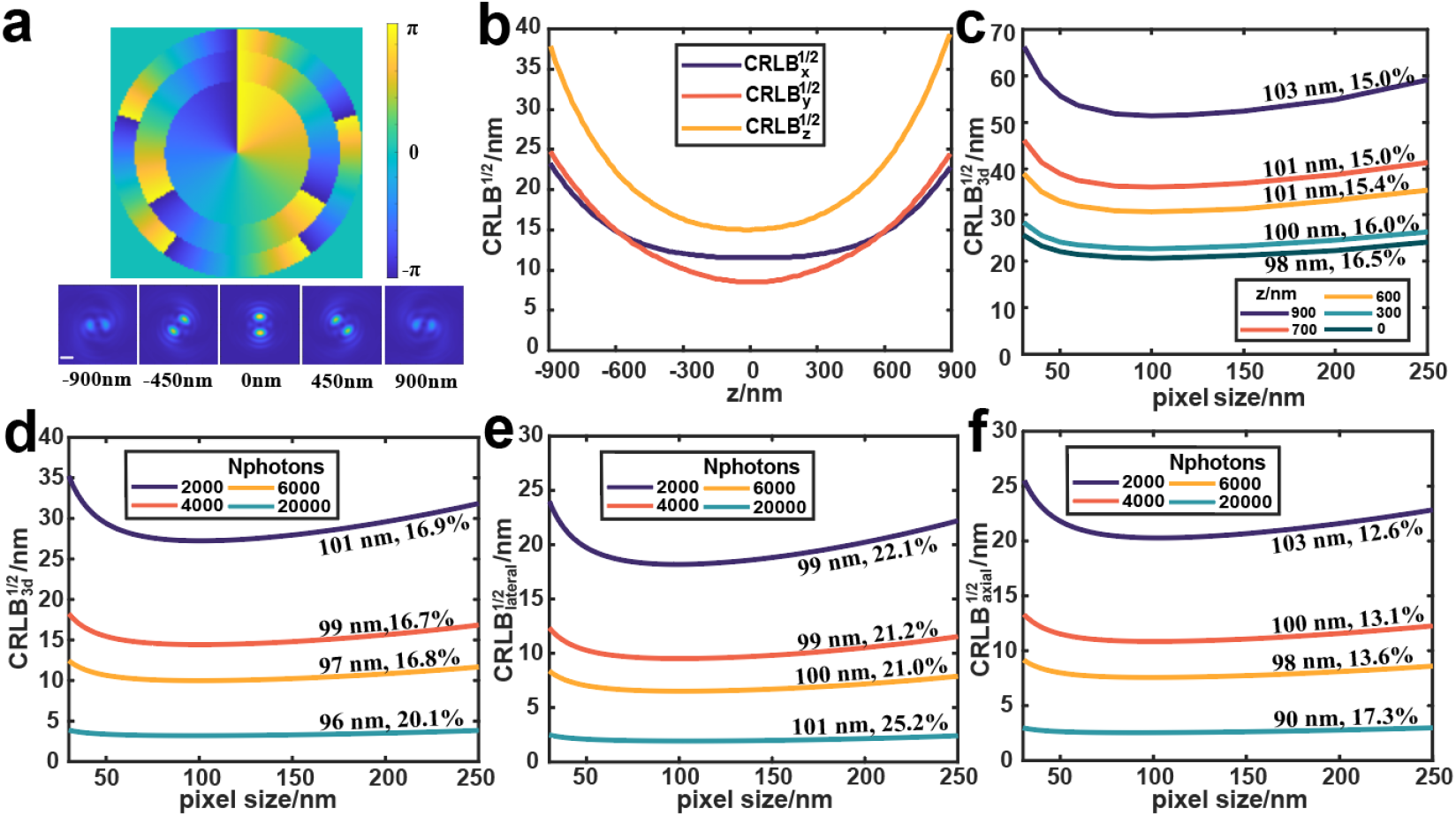
DH PSF simulation and CRLB calculation. (a) DH pupil and corresponding PSF. (b) 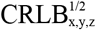 at different focal plane. (c) 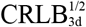 versus pixel size at different focal plane. (d-f) CRLB^½^ versus pixel size under different SBR, for 3d, lateral dimension, axial dimension, respectively. Scale bar 1*μ*m.

### 3.3. Optimal sampling rate for DMO Tetrapod PSF

Tetrapod PSF is a series of PSF which could achieve the best theoretical localization precision within a predefined axial range [18]. Here, we optimized a Tetrapod PSF with 6 *μ*m depth of field using the influence function of the deformable mirror actuators, which could achieve the practical best localization precision for a given deformable mirror [21]. The corresponding pupil function and 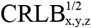 at different z of Tetrapod PSF are shown in Fig.4a and b. Next, we calculated the 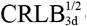 of Tetrapod PSF under different pixel size at different focal plane and found a more dramatic deterioration from 12.5% to 42.0% in CRLB sensitivity from defocus area (1500 nm above focal plane) to focal plane. The reason is probably that Tetrapod PSF near focus has finer patterns (4 main lobes) while the photons mainly localize in 2 lobes in the defocus area. Due to the complex Tetrapod PSF pattern, the optimal pixel size for 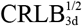 of Tetrapod PSF (89 nm) is relatively smaller compared to that of astigmatic PSF (93 nm) and DH-PSF (101 nm) for *N*_*phot*_ = *2*000. We then evaluated the effect of SBR on the optimal pixel size. As shown in Fig. 4d, the optimal pixel size for 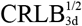 decreases from 89 nm to 81 nm as the increase of *N*_*phot*_ (from 2000 to 20000 photons). Under high SBR, 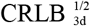 is also more sensitive to the pixelation effect. In Fig.4 e-f, it shows that the 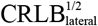 of Tetrapod PSF is more sensitive than 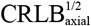 to the change of pixel size. Moreover, as *N*_*phot*_ increases from 2000 to 20000, optimal pixel size for 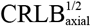 decreases from 95 nm to 77 nm, while optimal pixel size for 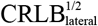 remains around 84nm.

**Fig. 4.**
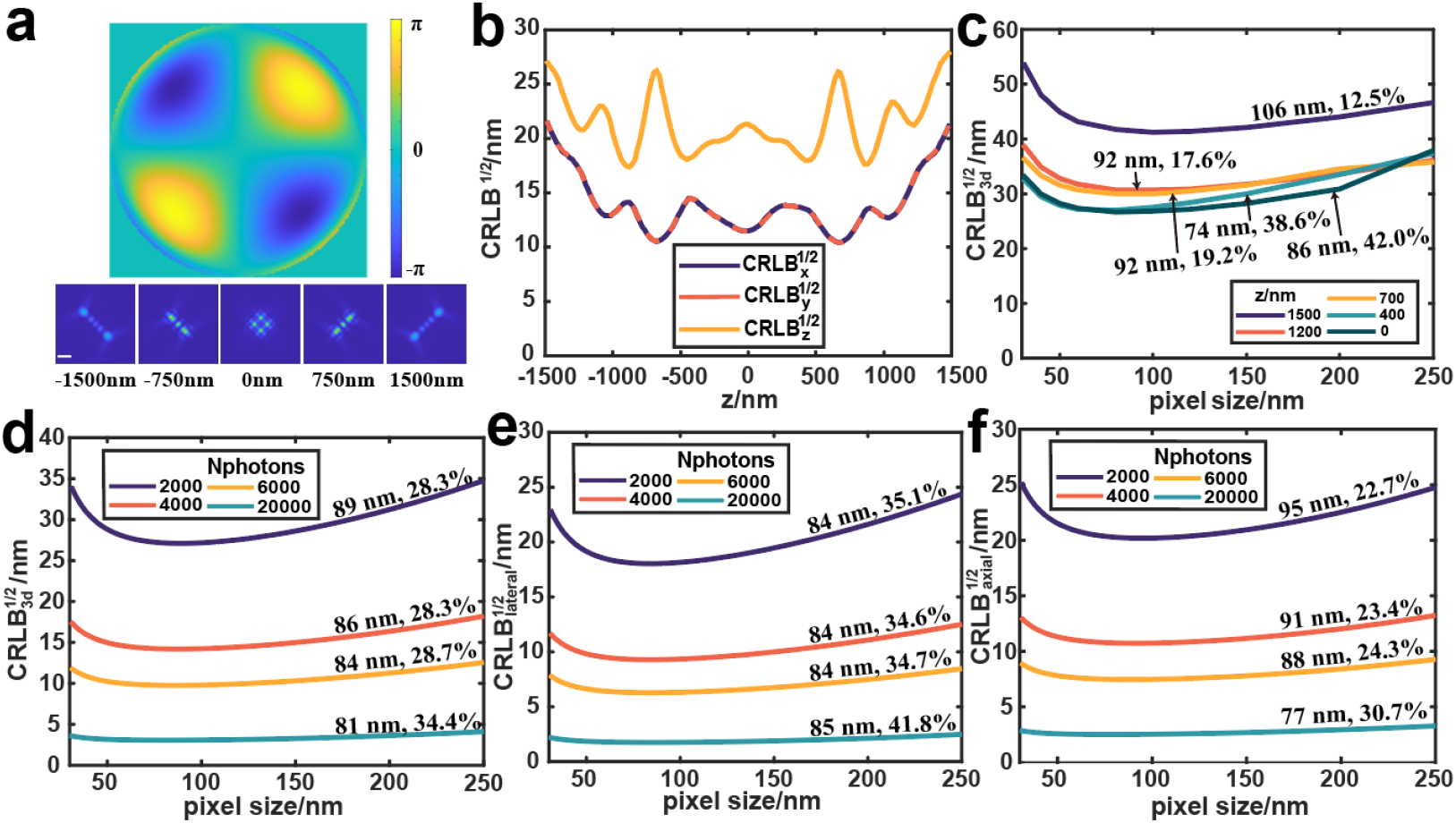
Tetrapod PSF simulation and CRLB calculation. (a) Tetrapod pupil and corresponding PSF, axial range analyzed is ±1500nm. (b) 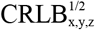 at different focal plane. (c) 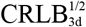 versus pixel size at different focal plane. (d-f) CRLB^½^ versus pixel size under different SBR, for 3d, lateral dimension, axial dimension, respectively. Scale bar 1 μm.

### 3.4. Optimal sampling rate for 4Pi PSF

4Pi microscope utilizes two opposing objectives to coherently detect the fluorescence signals and result in drastically increase of the axial resolution [26-28]. This approach for 3D localization is fundamentally different from the previously mentioned engineered PSF. The axial position is precisely determined by the interference phase, rather than the deformation of the PSF by defocus. Here, 4Pi PSF with four interference phases was used for evaluation. As expected, 4Pi-PSF showed the best 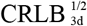 compared to the previously mentioned 3 engineered PSFs (Fig. 5b). Similarly, we first evaluate 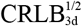 versus pixel size at different focal planes. Different from the other three PSFs, 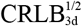 of 4Pi PSF has the lowest sensitivity to pixel size at focal plane (19.0%), while it exhibits strong sensitivity at defocus (43.8%, Fig. 5c). It is probably because 4Pi PSF has a concentrated pattern at focal plane, which contains less fine interference pattern compared to that at defocus positions. We next explored SBR influence. Here, the photon number of each objective of the 4Pi PSF was set the same as other PSFs, thus double photons for the entire 4Pi PSF. *N*_*phot*_ here denotes photons for each channel. As shown in Fig. 5d, the optimal sampling rate for *N*_*phot*_ = 1000 is about 97 nm. Under the same photon condition, 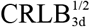 increases by ∼36.7% for 4Pi-PSFs with pixel size from 97 nm to 250 nm. It showed the strongest change among the four PSFs evaluated in this work. Furthermore, we systematically calculated the optimal pixel sizes of 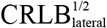 and 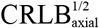 at different SBR levels. For *N*_*phot*_ from 1000 to 10000 photons, the optimal pixel size for 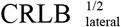 of different photons increases from 97 nm to 103 nm and the sensitivity decreases from 38.3% to 30.4%. For 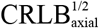, the optimal pixel size decreases from 87 nm to 74 nm and the sensitivity increases from 16.6% to 20.2%, whose trend is opposite compared to that of 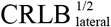.

**Fig. 5.**
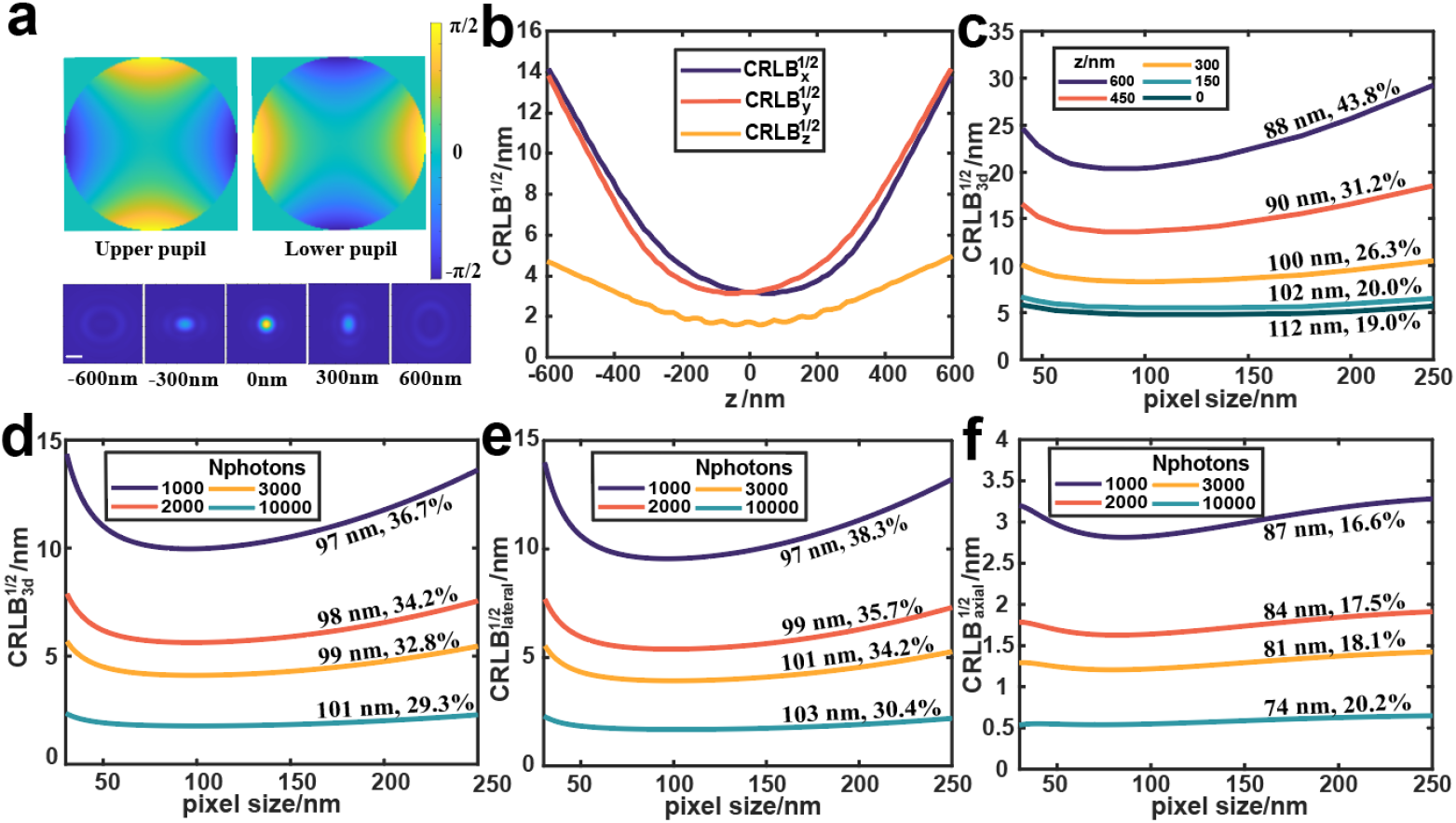
4Pi PSF simulation and CRLB calculation. (a) 4Pi pupil and corresponding PSF (interference phase is 0), axial range analyzed is ±600 nm. (b) 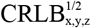 at different focal plane. (c) 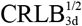 versus pixel size at different focal plane. (d-f) CRLB^½^ versus pixel size under different SBR, for 3d, lateral dimension, axial dimension, respectively. Scale bar 400 nm.

### 3.5 Potential effects from the sCMOS noise

sCMOS camera has become popular in scientific imaging for its large FOV and fast readout speed. However, pixel-dependent readout noise and hot pixel are also introduced to the signal. Hot pixel refers to the pixel with extremely high readout noise. As sampling rate changes, hot pixel effect may also exhibit differently. Therefore, we explored the hot pixel effect to CRLB at different sampling rates. We first simulated an astigmatic PSF at focus with fine sampling rates and binned it into PSFs with different pixel sizes. All the PSF model parameters are the same as described in the beginning of section 3, except for *N*_*phot*_ = *2*00, *bg* = 1 × 10^−4^ *photons/nm*^*2*^. We then selected a center pixel and gave 20 photoelectrons extra readout noise to simulate a hot pixel. We explored 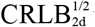 under different sampling rates for both uniform-readout-noise and hot-pixel-added conditions (Fig. 6). The results indicated that a single hot pixel only has tiny impact on the localization accuracy. The influence is a bit more obvious under a low sampling rate where each single molecule pattern contains fewer effective pixels.

**Fig. 6.**
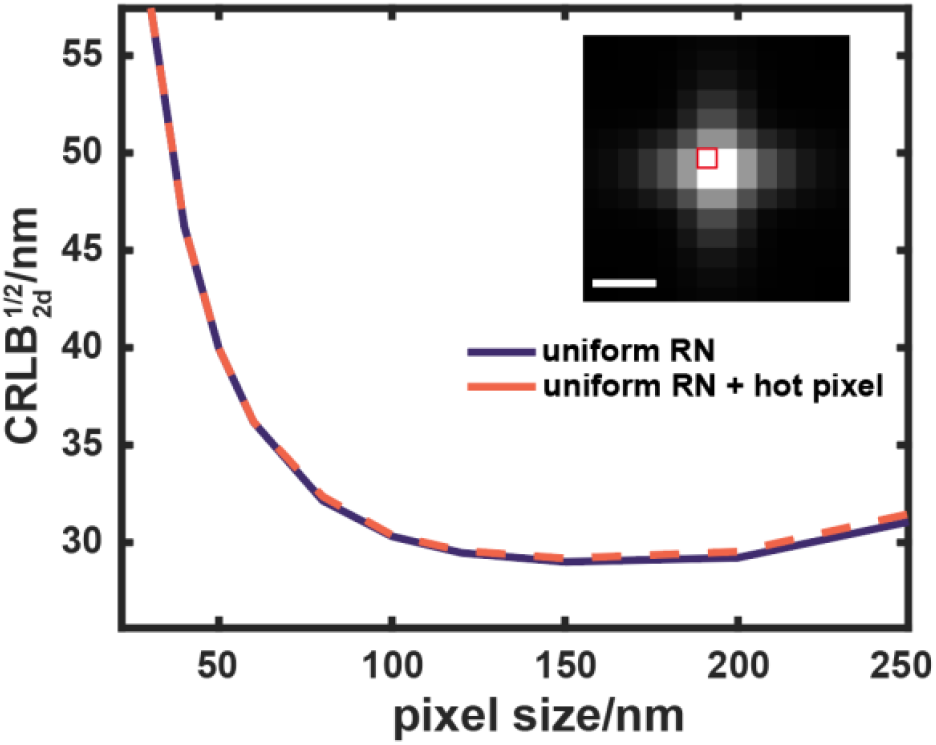
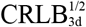 of astigmatic PSF by EMCCD with uniform readout noise(purple) and sCMOS with a hot pixel at the center(orange). The hot pixel is added with 20 photoelectrons extra readout noise and its position is marked with red frame. The pixel size of schematic PSF used in this figure is 100 nm. Scale bar 400 nm.

## 4. Discussion

The sampling rate of pixelated images is a critical factor for the precision of the single molecule localization. On the other hand, it also affects the FOV for a camera sensor chip with fixed number of pixels. However, this topic has not yet been studied in detail, especially for 3D single molecule localization. Based on the precise vector PSF model for high NA objective, we binned the fine sampled PSF models (2 nm), which accurately accounts for the pixel integration effect, to generate the pixelated PSFs. We systematically evaluate the effect of the pixel size on the achievable CRLB for 3D localization and CRLB sensitivity (defined as relative increase of CRLB^½^ from optimal sampling rate to 250 nm sampling rate) of four different PSFs at different defocus positions (Tab. 1). Astigmatic PSF and Tetrapod PSF show an increase in optimal sampling rate when defocusing, while 4Pi PSF shows the opposite effect with change of focal plane. The optimal sampling rate of DH PSF was almost the same for different focal positions. We then explored these PSFs’ optimal sampling rate and CRLB sensitivity under different SBR (Tab. 2). We found that PSFs with smaller features need finer sampling to achieve the optimal localization precision. For example, under 2,000 photons, optimal sampling rate for Tetrapod PSF which has abundant features is 89 nm, while it is 101 nm for DH PSF (Tab. 2). For a 4-channel 4Pi PSF, optimal sampling rate is 97 nm.

**Table 1.** Optimal sampling rate (OSR) and sensitivity of different PSFs at different z (Nphotons=2000, bg = 2 ×10^-3^/nm^2^)

| Astigmatic PSF |  |  | DH PSF |  |  | Tetrapod PSF |  |  | 4Pi PSF |  |  |
| --- | --- | --- | --- | --- | --- | --- | --- | --- | --- | --- | --- |
| z/<br>nm | OSR<br>/nm | sensi<br>tivity | z/<br>nm | OSR<br>/nm | sensi<br>tivity | z/<br>nm | OSR<br>/nm | sensi<br>tivity | z/<br>nm | OSR<br>/nm | sensi<br>tivity |
| 0 | 66 | 50.4% | 0 | 98 | 16.5% | 0 | 86 | 42.0% | 0 | 112 | 19.0% |
| 200 | 84 | 29.4% | 300 | 100 | 16.0% | 400 | 74 | 38.6% | 150 | 102 | 20.0% |
| 400 | 86 | 21.5% | 600 | 101 | 15.4% | 700 | 92 | 19.2% | 300 | 100 | 26.3% |
| 500 | 99 | 16.3% | 700 | 101 | 15.0% | 1200 | 92 | 17.6% | 450 | 90 | 31.2% |
| 600 | 105 | 12.5% | 900 | 103 | 15.0% | 1500 | 106 | 12.5% | 600 | 88 | 43.8% |

**Table 2.**
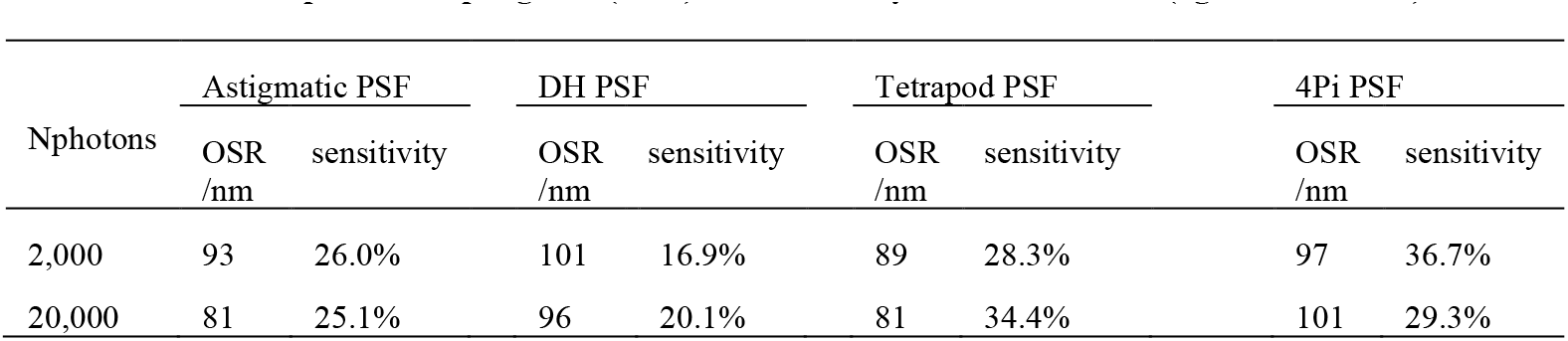
Optimal sampling rate (OSR) and sensitivity of different PSFs (bg = 2×10^-3^/nm^2^)

| Nphotons | Astigmatic PSF |  | DH PSF |  | Tetrapod PSF |  | 4Pi PSF |  |
| --- | --- | --- | --- | --- | --- | --- | --- | --- |
|  | OSR<br>/nm | sensitivity | OSR<br>/nm | sensitivity | OSR<br>/nm | sensitivity | OSR<br>/nm | sensitivity |
| 2,000 | 93 | 26.0% | 101 | 16.9% | 89 | 28.3% | 97 | 36.7% |
| 20,000 | 81 | 25.1% | 96 | 20.1% | 81 | 34.4% | 101 | 29.3% |

To intuitively showcase this intrinsic characteristic of PSF, we simulated astigmatic PSFs within axial range of ±100 nm around focus using different NA. A high-NA microscope collects more photons from the emitter with a larger opening angle, making the PSF sharper spatially. Consequently, such a PSF with more high frequency information (fine pattern) should exhibit a smaller optimal sampling rate based on the observation above. As shown in Fig. 7, we found that as the increase of NA from 1.0 to 1.5, optimal sampling rate of these PSFs drops from 76 nm to 60 nm, while their sensitivity to sampling rate increases from 25.2% to 54.0%, which is identical with our supposition.

**Fig. 7.**
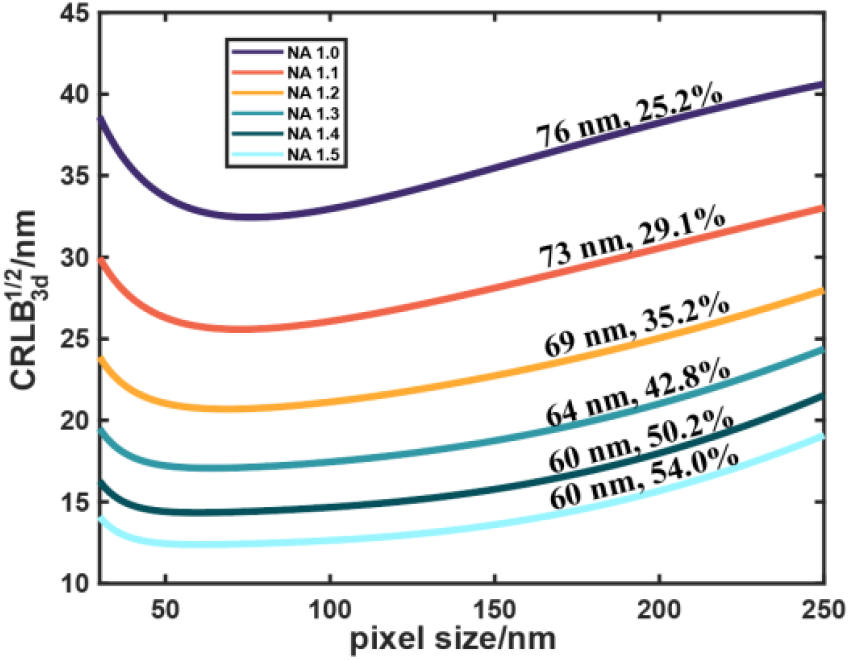
CRLB of an astigmatic PSF within ±100 nm axial range around focus using different NA for simulation. Each curve is marked with corresponding optimal sampling rate and sensitivity (defined as the relative increase of CRLB from optimal sampling rate to sampling rate of 250 nm).

Noise also affected the CRLB response to the optimal sampling rate. A fine sampling rate introduces too much read noise that deteriorates CRLB. When SBR increases, optimal pixel sizes for astigmatic PSF, DH PSF and Tetrapod PSF slightly decrease, while it remains almost unchanged for 4Pi PSF. Moreover, we found that the sCMOS hot pixels around single molecule have almost no influence on the optimal sample rate.

In summary, we adopted a binning method to simulate 4 different kinds of PSFs at different sampling rates and provide a framework to calculate CRLB of pixelated PSF with high accuracy. Our analysis pointed out that different PSFs have different amount of delicate features and this intrinsic characteristic accounts for their various optimal sampling rates and sensitivity. Moreover, the optimal sampling rates and sensitivity are also affected by SBR to a certain degree. We believe that the theoretical analysis performed in this work provides useful guide for the magnification design of optical systems in coordinate with PSF optimization to reach the best precision.

## Data availability

The code used in this work is available in Ref. [29].

## References

1. E. Betzig, G. H. Patterson, R. Sougrat, O. W. Lindwasser, S. Olenych, J. S. Bonifacino, M. W. Davidson, J. Lippincott-Schwartz, and H. F. Hess, “Imaging Intracellular Fluorescent Proteins at Nanometer Resolution,” Science 313(5793), 1642–1645 (2006).

2. S. T. Hess, T. P. K. Girirajan, and M. D. Mason, “Ultra-High Resolution Imaging by Fluorescence Photoactivation Localization Microscopy,” Biophysical Journal 91(11), 4258–4272 (2006).

3. M. J. Rust, M. Bates, and X. Zhuang, “Sub-diffraction-limit imaging by stochastic optical reconstruction microscopy (STORM),” Nature Methods 3(10), 793–796 (2006).

4. F. Balzarotti, Y. Eilers, K. C. Gwosch, A. H. Gynnå, V. Westphal, F. D. Stefani, J. Elf, and S. W. Hell, “Nanometer resolution imaging and tracking of fluorescent molecules with minimal photon fluxes,” Science 355(6325), 606–612 (2017).

5. C. S. Smith, N. Joseph, B. Rieger, and K. A. Lidke, “Fast, single-molecule localization that achieves theoretically minimum uncertainty,” Nature Methods 7(5), 373–375 (2010).

6. Y. Li, M. Mund, P. Hoess, J. Deschamps, U. Matti, B. Nijmeijer, V. J. Sabinina, J. Ellenberg, I. Schoen, and J. Ries, “Real-time 3D single-molecule localization using experimental point spread functions,” Nature Methods 15(5), 367–369 (2018).

7. D. R. L. Russell E. Thompson, Watt W. Webb, “Precise Nanometer Localization Analysis for Individual Fluorescent Probes,” Biophysical Journal 82(5), 9 (2002).

8. R. J. Ober, S. Ram, and E. S. Ward, “Localization Accuracy in Single-Molecule Microscopy,” Biophysical Journal 86(2), 1185–1200 (2004).

9. K. I. Mortensen, L. S. Churchman, J. A. Spudich, and H. Flyvbjerg, “Optimized localization analysis for single-molecule tracking and super-resolution microscopy,” Nature Methods 7(5), 377–381 (2010).

10. B. Huang, W. Wang, M. Bates, and X. Zhuang, “Three-Dimensional Super-Resolution Imaging by Stochastic Optical Reconstruction Microscopy,” Science 319(5864), 810–813 (2008).

11. S. R. P. Pavani, M. A. Thompson, J. S. Biteen, S. J. Lord, N. Liu, R. J. Twieg, R. Piestun, and W. E. Moerner, “Three-dimensional, single-molecule fluorescence imaging beyond the diffraction limit by using a double-helix point spread function,” Proceedings of the National Academy of Sciences 106(9), 2995–2999 (2009).

12. S. Prasad, “Rotating point spread function via pupil-phase engineering,” Opt. Lett. 38(4), 585–587 (2013).

13. M. Backlund, M. Lew, A. Backer, S. Sahl, G. Grover, A. Agrawal, R. Piestun, and W. Moerner, “The double-helix point spread function enables precise and accurate measurement of 3D single-molecule localization and orientation” Proc. SPIE 8590, 85900L (2013).

14. Y. Shechtman, S. J. Sahl, A. S. Backer, and W. E. Moerner, “Optimal Point Spread Function Design for 3D Imaging,” Physical Review Letters 113(13), 133902 (2014).

15. S. Hell, and E. H. K. Stelzer, “Properties of a 4Pi confocal fluorescence microscope,” J. Opt. Soc. Am. A 9(12), 2159–2166 (1992).

16. K. Bahlmann, S. Jakobs, and S. W. Hell, “4Pi-confocal microscopy of live cells,” Ultramicroscopy 87(3), 155–164 (2001).

17. H. Gugel, J. Bewersdorf, S. Jakobs, J. Engelhardt, R. Storz, and S. W. Hell, “Cooperative 4Pi Excitation and Detection Yields Sevenfold Sharper Optical Sections in Live-Cell Microscopy,” Biophysical Journal 87(6), 4146–4152 (2004).

18. Y. Shechtman, L. E. Weiss, A. S. Backer, S. J. Sahl, and W. E. Moerner, “Precise Three-Dimensional Scan-Free Multiple-Particle Tracking over Large Axial Ranges with Tetrapod Point Spread Functions,” Nano Lett 15(6), 4194–4199 (2015).

19. S. Stallinga, and B. Rieger, “Accuracy of the Gaussian Point Spread Function model in 2D localization microscopy,” Opt. Express 18(24), 24461–24476 (2010).

20. M. E. Siemons, L. C. Kapitein, and S. Stallinga, “Axial accuracy in localization microscopy with 3D point spread function engineering,” Opt. Express 30(16), 28290–28300 (2022).

21. S. Fu, M. Li, L. Zhou, Y. He, X. Liu, X. Hao, and Y. Li, “Deformable mirror based optimal PSF engineering for 3D super-resolution imaging,” Opt. Lett. 47(12), 3031–3034 (2022).

22. M. Leutenegger, R. Rao, R. A. Leitgeb, and T. Lasser, “Fast focus field calculations,” Opt. Express 14(23), 11277–11291 (2006).

23. A. Abraham, S. Ram, J. Chao, E. S. Ward, and R. Ober, “Comparison of estimation algorithms in single-molecule localization” Proc. SPIE 7570, 757004 (2010).

24. F. Huang, T. M. P. Hartwich, F. E. Rivera-Molina, Y. Lin, W. C. Duim, J. J. Long, P. D. Uchil, J. R. Myers, M. A. Baird, W. Mothes, M. W. Davidson, D. Toomre, and J. Bewersdorf, “Video-rate nanoscopy using sCMOS camera–specific single-molecule localization algorithms,” Nature Methods 10(7), 653–658 (2013).

25. C. Roider, A. Jesacher, S. Bernet, and M. Ritsch-Marte, “Axial super-localisation using rotating point spread functions shaped by polarisation-dependent phase modulation,” Opt. Express 22(4), 4029–4037 (2014).

26. Y. Li, E. Buglakova, Y. Zhang, J. V. Thevathasan, J. Bewersdorf, and J. Ries, “Accurate 4Pi single-molecule localization using an experimental PSF model,” Opt. Lett. 45(13), 3765–3768 (2020).

27. J. Wang, E. S. Allgeyer, G. Sirinakis, Y. Zhang, K. Hu, M. D. Lessard, Y. Li, R. Diekmann, M. A. Phillips, I. M. Dobbie, J. Ries, M. J. Booth, and J. Bewersdorf, “Implementation of a 4Pi-SMS super-resolution microscope,” Nature Protocols 16(2), 677–727 (2021).

28. J. Chen, B. Yao, Z. Yang, W. Shi, T. Luo, P. Xi, D. Jin, and Y. Li, “Ratiometric 4Pi single-molecule localization with optimal resolution and color assignment,” Opt. Lett. 47(2), 325–328 (2022).

29. H. Chang, S. Fu and Y. Li “Optimal Sampling Rate for Single Molecule Localization,” GitHub (2023) [accessed 13 September 2023] https://github.com/Li-Lab-SUSTech/Optimal-Sampling-Rate-of-Different-PSFs-in-SMLM.

